# Schedule-dependent production of stereotyped sequences of actions

**DOI:** 10.1101/2022.06.16.496442

**Authors:** Emma G. Follman, Maxime Chevée, Courtney J. Kim, Amy R. Johnson, Jennifer Tat, Erin S. Calipari

## Abstract

Concatenating actions into automatic routines is evolutionarily advantageous as it allows organisms to efficiently use time and energy under predictable conditions. However, over reliance on inflexible behaviors can be life-threatening in a changing environment and can become pathological in disease states such as obsessive-compulsive disorder (OCD) and substance use disorder (SUD). Understanding the conditions under which stereotypical sequences of actions are produced is crucial to studying how these behaviors can become maladaptive. Here, we investigated the ability of operant conditioning schedules and contingencies to promote reproducible sequences of five lever presses. We found that signaling reinforcer delivery with a visual cue was effective at increasing learning rates but resulted in mice pressing the lever in fast succession until the cue turned on, rather than pressing it five times. We also found that requiring mice to collect their reinforcer between sequences had little effect on both rate of behavior and on quantitative metrics of reproducibility such as interresponse interval (IRI) variance, and that a training strategy that directly reinforced sequences with low variance IRIs was not more effective than a traditional fixed ratio schedule at promoting reproducible action execution. Together, our findings provide insights into the parameters of behavioral training that promote reproducible sequences and serve as a roadmap to investigating the neural substrates of automatic behaviors.

## Introduction

The ability to learn relationships between actions and outcomes is the basis of reinforcement behavior. Early stages of reinforcement learning are probabilistic - initially a random action results in an outcome and then that action is more likely or less likely to occur in the future, depending on the consequence (Gershman and Ölveczky, 2020). This process requires a large amount of unpredictable engagement with the environment as the organism learns the contingencies and relationships of the task; however, over time these relationships can become more predictable. As a behavior is practiced even more, it begins to become automatic (Dezfouli and Balleine, 2012). Habitual, automatic behaviors allow us to perform routine actions without wasting time and energy on decision making (Graybiel, 2008). However, they are also less sensitive to changes in consequences, which can lead to sustaining unhealthy behaviors that can degenerate into pathological conditions such as obsessivecompulsive disorder (Burguière et al., 2015) and addiction (Pierce and Vanderschuren, 2010).

How automatic sequences of actions are learned and what factors contribute to their development is not well understood. While the study of human behaviors both in health (Keller et al., 2021; Luque et al., 2020; Smith and Graybiel, 2016) and disease (Burguière et al., 2015; Byrne et al., 2021; O’Tousa and Grahame, 2014) provides useful insights into these mechanisms, many investigations of the molecular and circuit underpinnings of automatic behaviors are currently performed in rodents (Bouton, 2021; Burguière et al., 2015; Faure et al., 2005; Gremel and Costa, 2013; Lerner, 2020; Renteria et al., 2018; Wassum et al., 2009) due to the availability of genetic and optogenetic tools. Understanding the features of training paradigms that effectively or ineffectively promote the emergence of reproducible behavioral sequences is therefore necessary to take full advantage of the tools only accessible to rodent research, and to identify the circuits, cell types and molecular mechanisms that control sequences of stereotypical actions.

In this study, we trained mice in a series of operant conditioning tasks where sucrose delivery was reinforced on a fixed ratio 5 (FR5) schedule of reinforcement. We tested three distinct hypotheses: First, we tested whether the presence of an additional cue presented concurrently with sucrose delivery influenced the pattern of lever pressing throughout learning. A “signaled” reward has been shown to improve learning (Branch, 1977; Lewis et al., 1974; Marcucella & Margolius, 1978; Sanderson et al., 2014; Schachtman & Reed, 1992), thus we reasoned that the immediate feedback provided by the cue and the absence of the need to check for reinforcer delivery after completing a sequence of actions may promote sequence termination and the development of more precise sequences. Second, we tested whether requiring animals to collect their earned sucrose, thus preventing them from accumulating unconsumed reinforcers, was an effective condition to promote the generation of sequences. Third, and finally, we tested whether reinforcing reproducible sequences by only rewarding sequences whose inter-response interval (IRI) variance was below a target was an effective strategy to promote stereotypical patterns of operant behavior.

Our results show that indeed, signaling reinforcer delivery with a consequent cue is crucial for mice to learn and to develop sequencing behavior. The sucrose collection condition, however, was superfluous regarding most metrics quantifying reproducibility of behaviors. Finally, the strategy in which we directly reinforced reproducible sequences revealed that signaling reinforcer delivery with a cue, while very effective at promoting learning and sequencing behavior, promotes lever pressing in bouts terminated by stimulus presentation rather than sequences characterized by their intrinsic number of presses. Our results provide valuable insight into the features that drive sequential behaviors and will allow future studies to investigate the neural substrates of such behaviors while carefully tuning their training parameters.

## Methods

### Subjects

Experiments were approved by the Institutional Animal Care and Use Committee of Vanderbilt University Medical Center and conducted according to the National Institutes of Health guidelines for animal care and use. Forty-seven 8-week-old animals were used for this study. C57BL/6J mice (22 males and 25 females) were acquired from Jackson Laboratory (Bar Harbor, ME; SN: 000664) and maintained on an 8am/8pm 12-hour reverse light cycle. Experiments were performed during the dark phase. Four to five animals were housed per cage with unlimited access to water. Food access was restricted to maintain ~90% pre-restriction body weight. Only mice that met FR1 acquisition criteria (as described below) were moved to each subsequent FR5 sessions.

The number of animals in each group is as follows: 1 *FR5 w/MustCollect* mouse did not reach FR1 acquisition criteria and was excluded. An additional 9 mice were lost before completing 10 days of FR5 and were also excluded from all subsequent analyses (N = 1 from the *FR5 w/ MustCollect* group, N = 6 from the *FR5 w/LightCue&MustCollect* group and N = 2 from the *LowVariance* group). In total, the cohort reported in this study included 37 animals (*FR5 w/LightCue&MustCollect:* 5 males/4 females; *FR5 w/ MustCollect*: 3 males/3 females; *FR5 w/LightCue*: 4 males/4 females*; LowVariance*: 5 males/9 females). The data in Figure 5 were only acquired for a subset of animals (N=9 mice from the *LowVariance* group, N=6 mice from the *FR5 w/ LightCue&MustCollect* group).

### Apparatus

Mice were trained and tested daily in individual standard wide mouse operant conditioning chambers (Med Associates Inc., St. Albans, Vermont) in which 3D-printed dividers were inserted, limiting the available space to a small square area providing access to a sucrose port and a lever (area: 13×13=169cm^2^). These boxes were fitted with a standard retractable lever and a white noise generator with a speaker. A custom-made 3D-printed wall insert was used to hold and display a stainless-steel cannula (lick port, 18 gauge, 0.042 “ID, 0.05” OD, 0.004” Wall Thickness), which was connected to a syringe pump for sucrose delivery. An illumination light was affixed above the lick port. To measure lever displacement, two small magnets (Neodymium block magnets N45 0.069×0.591×0.197 in, Buymagnets.com #EP331) were fixed to the lever and a Hall effect sensor (Sensor Hall Analog Radial Lead, Honeywell #SS49E) was placed 5mm above the lever. The weight of the magnets was countered by attaching a 1.38” 12V 44LB electromagnet (APW company #EM137-12-222) 30mm above the lever. Control of the electromagnet strength and acquisition of the Hall effect sensor data were performed using an Arduino Nano Every (Arduino, #ABX00033).

### Procedure

#### General Procedural Information

All sessions lasted until the maximum number of rewards was obtained (51) or 1 hour was reached, whichever came first. White noise signaled the beginning of the session and was on for the entire duration of the session.

#### Task Design

##### FR5 w/MustCollect

Mice were first trained on a fixed ratio 1 (FR1) schedule of reinforcement. Each lever press resulted in the delivery of 8 μL of a 10% sucrose solution. Additional presses performed after sucrose delivery but before sucrose collection had no programmed consequence and did not count toward the next sequence. Once the reinforcer was collected, lever presses counted again. Acquisition criteria were considered met once a mouse had obtained 50 rewards within the allotted 1 hour for two consecutive days (5.7±2.0 days, N=3 females; 5.0±0.58 days, N=3 males). Mice that did not meet the criteria within 10 days were excluded (N=1 mouse). Upon meeting the criteria, the reinforcement contingency was increased to FR5 with all other conditions the same. All animals in the study were trained on this paradigm for 10 days.

##### FR5 w/LightCue

Animals were first trained on an FR1 schedule of reinforcement. Each lever press resulted in the delivery of 8 μL of a 10% sucrose solution and the light above the lick port turning on for 1 second. Acquisition criteria were considered met once a mouse had obtained 50 rewards within the allotted 1 hour for two consecutive days (3.25±0.63 days, N=4 females; 3.5±0.65 days, N=4 males). Mice that did not meet criteria within 10 days were excluded (N=0 mouse). Upon meeting the criteria, the reinforcement contingency was increased to FR5 with all other conditions the same. All animals in the study were trained on this paradigm for 10 days.

##### FR5 w/LightCue&MustCollect

Mice trained on this task experienced both conditions described above (*w/LightCue* and *w/MustCollect*). Acquisition criteria were considered met once a mouse had obtained 50 rewards within the allotted 1 hour for two consecutive days (3.7±0.33 days, N=3 females; 3.0±0.26 days, N=6 males). Mice that did not meet the criteria within 10 days were excluded (N=0 mouse). Upon meeting criteria, the reinforcement contingency was increased to FR5 with all other conditions the same. All animals in the study were trained on this paradigm for 10 days.

##### LowVariance

This strategy was identical to the *FR5 w/LightCue&MustCollect* group during the initial FR1 training phase, in that it included both the *w/ LightCue* and *w/ MustCollect* conditions (days to acquisitions: 4.1±0.42 days, N=9 females; 3.8±0.49 days, N=5 males). Subsequently, during the FR5 phase, a sequence of 5 presses was only rewarded if the variance of its within-sequence inter-response intervals (IRIs) was below a threshold computed as the median IRI variance over the last 5 sequences. If the IRI variance of a sequence was above the threshold at that time, the threshold was updated but no external signals were generated, and the animal simply had to try again.

### Analysis

#### Statistical Analyses

All analyses were performed using custom code in Python (v3.6.13). The SciPy package (v1.5.3) was used to perform paired t-tests (Figures 2A, 3F, 3G, 4D, 4E, 6B, 6C) and the Pingouin package (v0.3.12) was used to perform one-way and mixed ANOVAs as well as the corresponding post-hoc Tukey tests (Figures 2B, 2C, 2D, 3F, 3G, 5C). All data are reported as Mean ± SEM and all statistical tests used are specified in the Results section.

#### Hall effect sensor data processing

Data was acquired at 1,000 Hz and lowpass filtered at 5e^-9^ cycles/unit to remove fast oscillations originating from the electromagnet. To identify deflections in the time series, we first computed a threshold, which, when used to define deflection points, resulted in the same number of deflections as lever presses counted by MedPC. However, this procedure resulted in a small number of false positives as well as false negatives. To identify which deflections in the data represented a lever press counted by MedPC versus deflections that were too small to trigger a bonafide lever press, we used MedPC timestamps as a reference. This approach allowed us to manually adjust the labels for each deflection, and only sessions in which 90% of presses were accounted for were kept for further analysis. The start and end of each deflection were identified by sliding backwards and forwards in time, respectively, from the threshold crossing point until the values returned to baseline or until the preceding/following press. The data from each session was z-scored. To compute the Pearson correlation coefficient between individual presses, we used one second from the start of each deflection. Presses that lasted longer were truncated and presses that were shorted were padded with zeros. This approach resulted in the correlations being primarily driven by the shape of the downward deflection and the duration of the press. Changing the size of this window had little effect on our results. This system was only installed half-way through the experiment, which is why we only presented data for a subset of mice and days. Hall effect sensor analysis were performed with *Python 3.9.7*.

## Results

### Paradigm-specific effects on reinforcement behavior

To test the hypothesis that specific features of operant conditioning contribute to generating sequence behaviors, we trained four groups of mice on four distinct FR5 lever pressing procedures. A first group was trained on an FR5 schedule in which lever presses only counted if executed after the previous reinforcer had been collected (*FR5 w/MustCollect*, N=6, **Figure 1Ai**). A second group was trained on an FR5 schedule in which a light cue signaled sucrose delivery (one second light cue, *FR5 w/LightCue*, N=8, **Figure 1Aii**). A third group was trained with both conditions (*FR5 w/ LightCue&MustCollect*, N=9, **Figure 1Aiii**). Finally, a fourth group was trained to test whether a schedule that specifically reinforces stereotypical sequences of lever presses would be more effective than FR5 at producing highly reproducible actions. Specifically, the variance of the IRIs in a sequence of five presses had to be below a target variance to trigger reinforcer delivery. The target was dynamically defined as the median variance of the last 5 sequences (*LowVariance*, N=14, **Figure 1Aiv**). In the case of a failed sequence, the mouse did not get any indication that the IRI variance was higher than the target and simply had to continue pressing until it generated a sequence whose IRI variance was below the target. These four strategies allowed us to determine which specific features of training paradigms promote or hinder the development of reproducible sequences of actions.

**Figure 1:**
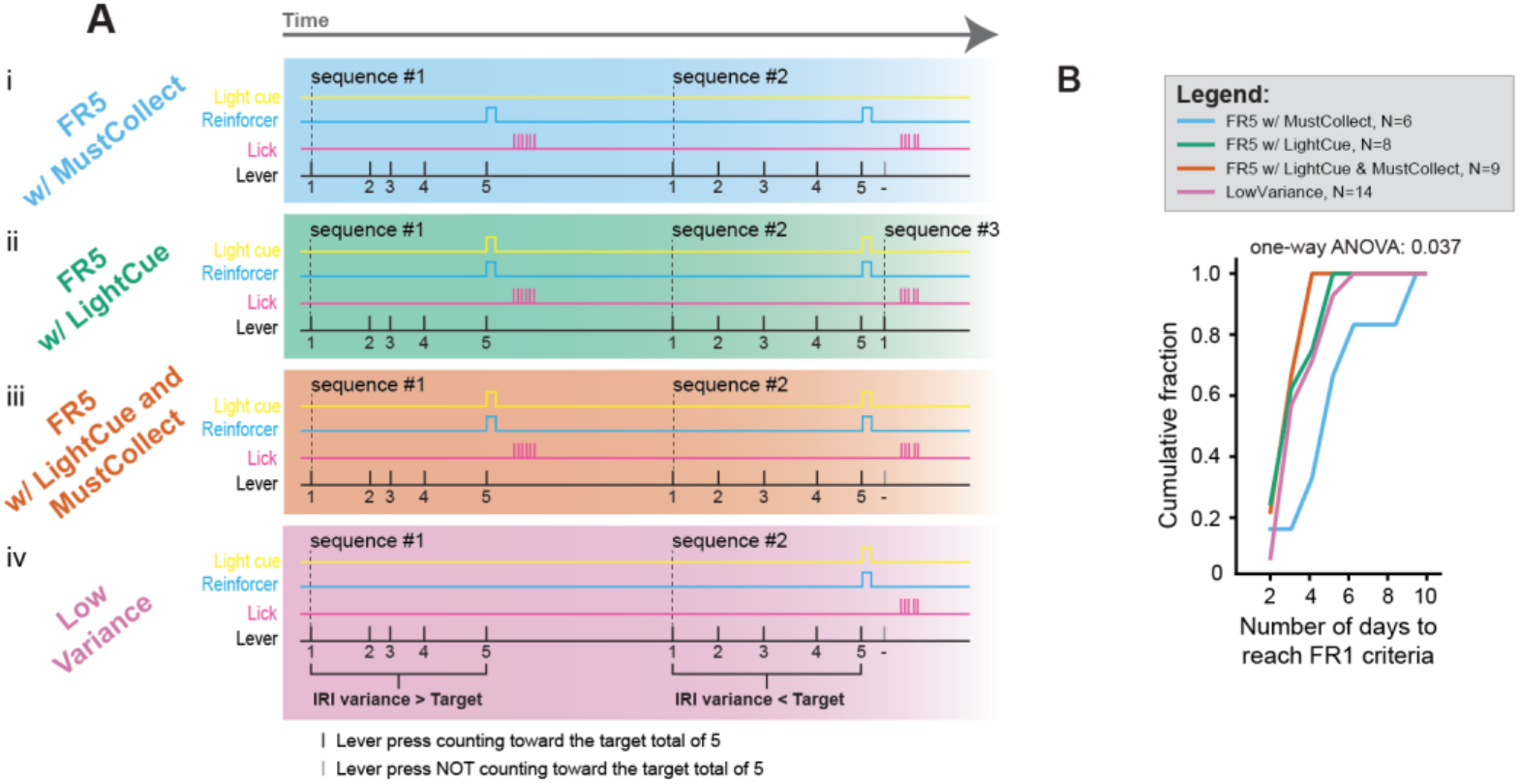
A light cue signaling sucrose delivery improves acquisition of a lever pressing task. **(A)** Diagram describing the four reinforcement strategies used in this study. **(B)** Cumulative fraction plot showing the number of days mice from each group took to reach the FR1 acquisition criteria (maximum rewards acquired on two consecutive days). Data presented as cumulative fraction of total mice.

### A light cue that signals sucrose delivery improves acquisition of a lever pressing operant task

All mice were first trained on an FR1 schedule and moved on to FR5 once they acquired the maximum number of reinforcers on two consecutive days (51 rewards, 1-hour max sessions). To determine whether the *w/ LightCue* and the *w/ MustCollect* conditions impacted the mice’s ability to learn to press the lever for a sucrose reinforcer, we plotted the cumulative distribution of days to FR1 acquisition for each group (**Figure 1B**). There was a significant difference in the time to acquisition across groups for mice that reached criteria within 10 days (*FR5 w/LightCue&MustCollect*, N=9 mice: 3.1±0.26 days; *FR5 w/MustCollect*, N=6 mice: 5.2±0.94 days; *FR5 w/LightCue*, N=8 mice: 3.4±0.42 days; *LowVariance*, N=14 mice: 3.7±0.30 days; one-way ANOVA, F = 3.2, p = 0.037), with the group trained using the *FR5 w/MustCollect* paradigm acquiring slower than the *FR5 w/ LightCue&MustCollect* group (Post-hoc Tukey test: *FR5 w/LightCue&MustCollect* _vs_ *FR5 w/MustCollect* p = 0.03). This result suggests that the light cue signaling reinforcer delivery improved the acquisition of a simple lever pressing task, a result in line with previous studies (Branch, 1977; Lewis et al., 1974).

### Absence of a light cue signaling reinforcer delivery impairs the response rate on an FR5 schedule of reinforcement

We first tested whether each training strategy successfully increased the rate of lever pressing across 10 sessions. Mice from all groups except *FR5 w/ MustCollect* pressed the lever at higher rates late in training (day 9/10) compared to early in training (day 1) (**Figure 2A,** *FR5 w/LightCue&MustCollect* - early: 5.7±0.42 LP/min, late: 11.8±1.3 LP/min, paired t-test p = 0.032; *FR5 w/MustCollect* - early: 2.2±0.18 LP/min, late: 3.4±0.58 LP/min, paired t-test p = 0.27; *FR5 w/LightCue* - early: 4.3±0.53 LP/min, late: 7.6±1.3 LP/min, paired t-test p = 0.036; *LowVariance* - early: 2.3±0.25 LP/min, late: 5.7±0.41 LP/min, paired t-test p = 4.1e^-5^). To identify which training strategy resulted in the largest increase in response rate, we computed the fold change in rate between late and early across groups that had a significant increase and found no difference across contingencies (**Figure 2B**, *FR5 w/ LightCue&MustCollect*: 2.6±0.70; *FR5 w/LightCue*: 1.8±0.32; *LowVariance*: 3.1±0.47; one-way ANOVA, F = 1.457, p = 0.25). However, the *FR5 w/LightCue&MustCollect* group pressed faster than *FR5 w/ MustCollect* and *LowVariance* late in training (**Figure 2C**, *FR5 w/ LightCue&MustCollect*: 11.8±1.8 LP/min; *FR5 w/MustCollect*: 3.4±0.86 LP/min; *FR5 w/ LightCue*: 7.6±1.8 LP/min; *LowVariance*: 5.7±0.59 LP/min; one-way ANOVA, F = 8.9, p = 4.3e-5; Posthoc Tukey tests: *FR5 w/LightCue&MustCollect* _vs_ *FR5 w/MustCollect* p = 0.0010, *FR5 w/LightCue&MustCollect _vs_LowVariance* p = 0.0010) and *FR5 w/LightCue&MustCollect* and *FR5 w/ LightCue* groups obtained more total rewards than the *FR5 w/ MustCollect* and *LowVariance* groups across sessions (**Figure 2D,** *FR5 w/LightCue&MustCollect*, N=15 mice:493±5.57 reinforcers; *FR5 w/MustCollect*, N=6 mice: 296±32.7 reinforcers; *FR5 w/ LightCue*, N=8 mice: 427±32.1 reinforcers; *LowVariance*, N=16 mice: 285±23.2 reinforcers; oneway ANOVA, F = 17.08, p = 7.17e^-7^; posthoc Tukey test: *FR5 w/LightCue&MustCollect* _vs_ *FR5 w/ MustCollect* p = 0.0010; *FR5 w/LightCue&MustCollect _vs_LowVariance* p = 0.0010; *FR5 w/ LightCue* _vs_ *FR5 w/ MustCollect* p = 0.016; *FR5 w/ LightCue _vs_LowVariance* p = 0.0010). These results indicate that all strategies except *FR5 w/MustCollect*, which was the only paradigm without a light cue signaling sucrose delivery, reinforced lever pressing, although they resulted in distinct rates of reinforcer delivery.

**Figure 2:**
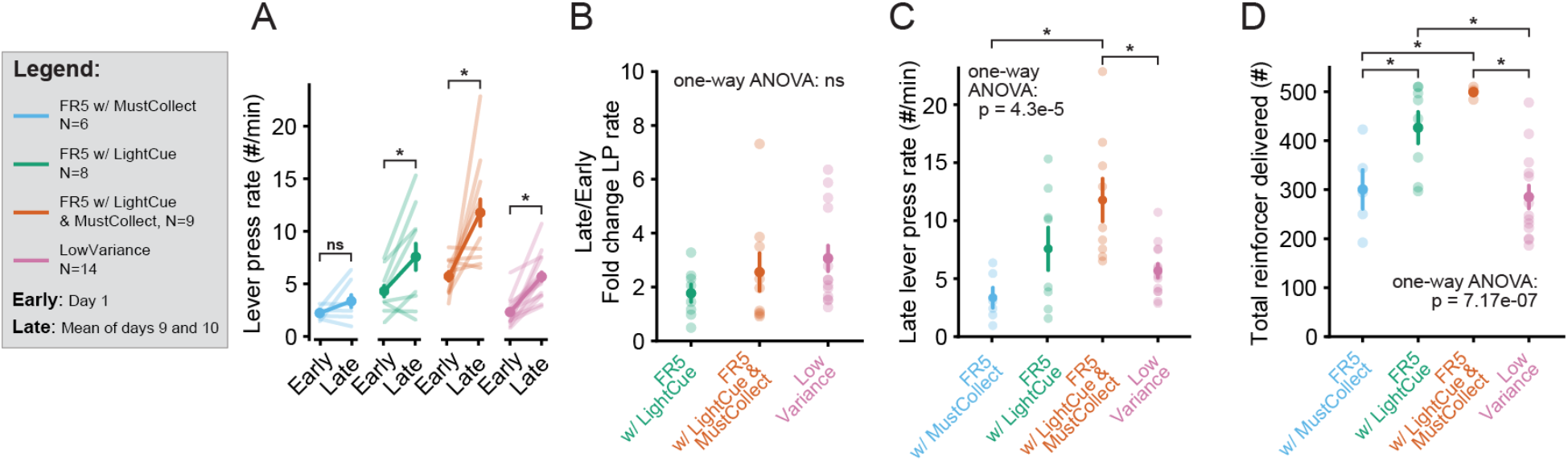
Absence of a light cue signaling sucrose delivery impairs performance on an FR5 lever pressing task. **(A)** Comparison of lever press rates between early in training (day 1) and late in training (mean of days 9 and 10) for each group. **(B)** Comparison of the fold change in lever press rates across groups which had a significant change. **(C)** Comparison of the lever press rates late in training across groups. **(D)** Comparison of the total rewards acquired during 10 days across groups. Data presented as mean +/- S.E.M. * p < 0.05, ns, not significant.

### A light cue that signals sucrose delivery is necessary for clustering lever presses into bouts

To determine how effective each training strategy was at promoting the clustering of presses into bouts, we analyzed the distribution of IRIs. We labeled IRIs between presses occurring within a reinforced sequence of five as “within-sequence IRIs” and IRIs between presses occurring across two subsequent sequences as “between-sequence IRIs” (**Figure 3A**). Mice that learn to perform a sequence of 5 presses should have “within-sequence IRIs” that are progressively shorter compared to “between-sequence IRIs”. To visualize this, we plotted a histogram showing the distribution of “within-sequence IRIs” and of “between-sequence IRIs” for the first and last sessions for one example mouse from each group, as well as heatmaps showing the same distributions across all sessions (**Figure 3B-E**). The median “within-sequence IRIs” decreased from early (day 1) to late (mean of days 9 and 10) in training for each group except for mice trained on the *FR5 w/ MustCollect* paradigm (**Figure 3F,** left, *FR5 w/ LightCue&MustCollect* - early: 2.8±0.24 s, late: 0.78±0.065 s; paired t-test p = 6.3e^-4^; *FR5 w/ MustCollect* - early: 9.1±1.1 s, late: 11.6±2.9 s; paired t-test p = 0.46; *FR5 w/LightCue* - early: 5.9±1.2 s, late: 1.3±0.21 s; paired t-test p = 0.035; *LowVariance* - early: 11.8±1.7 s, late: 1.5±0.14 s; paired t-test p = 7.5e^-4^), although there was no difference in the late/early fold change in “within-sequence IRI” duration between the three groups that clustered their lever presses (**Figure 3F,** right, *FR5 w/LightCue&MustCollect*: 0.32±0.055, *FR5 w/LightCue*: 0.36±0.90, *LowVariance*: 0.17±0.029; one-way ANOVA, F = 3.076, p =0.062). Similarly, the ratio of median “between-sequence IRIs” to median “within-sequence IRIs” increased for all groups except *FR5 w/MustCollect* (**Figure 3G**, left, *FR5 w/LightCue&MustCollect* - early: 3.3±0.27, late: 12.6±1.4; paired t-test p = 0.0022; *FR5 w/MustCollect* - early: 2.7±0.48, late: 3.4±1.0; paired t-test p = 0.71; *FR5 w/LightCue* - early: 2.9±0.29, late: 16.2±2.0; paired t-test p = 0.0036; *LowVariance* - early: 2.1±0.19, late: 19.2±3.1; paired t-test p = 0.0017), showing that all conditions but the one without a light cue signaling sucrose delivery clustered their lever presses into bouts. To compare the magnitude of the clustering attained with each training strategy, we compared the between/within ratio for the three groups which clustered their bouts and found no differences (**Figure 3G**, right, *FR5 w/LightCue&MustCollect*: 4.2±0.86, *FR5 w/ LightCue*: 6.8±2.0, *LowVariance*: 9.7±2.1; one-way ANOVA, F = 2.226, p =0.127), suggesting all three strategies promoted the clustering of lever presses into bouts equally.

**Figure 3:**
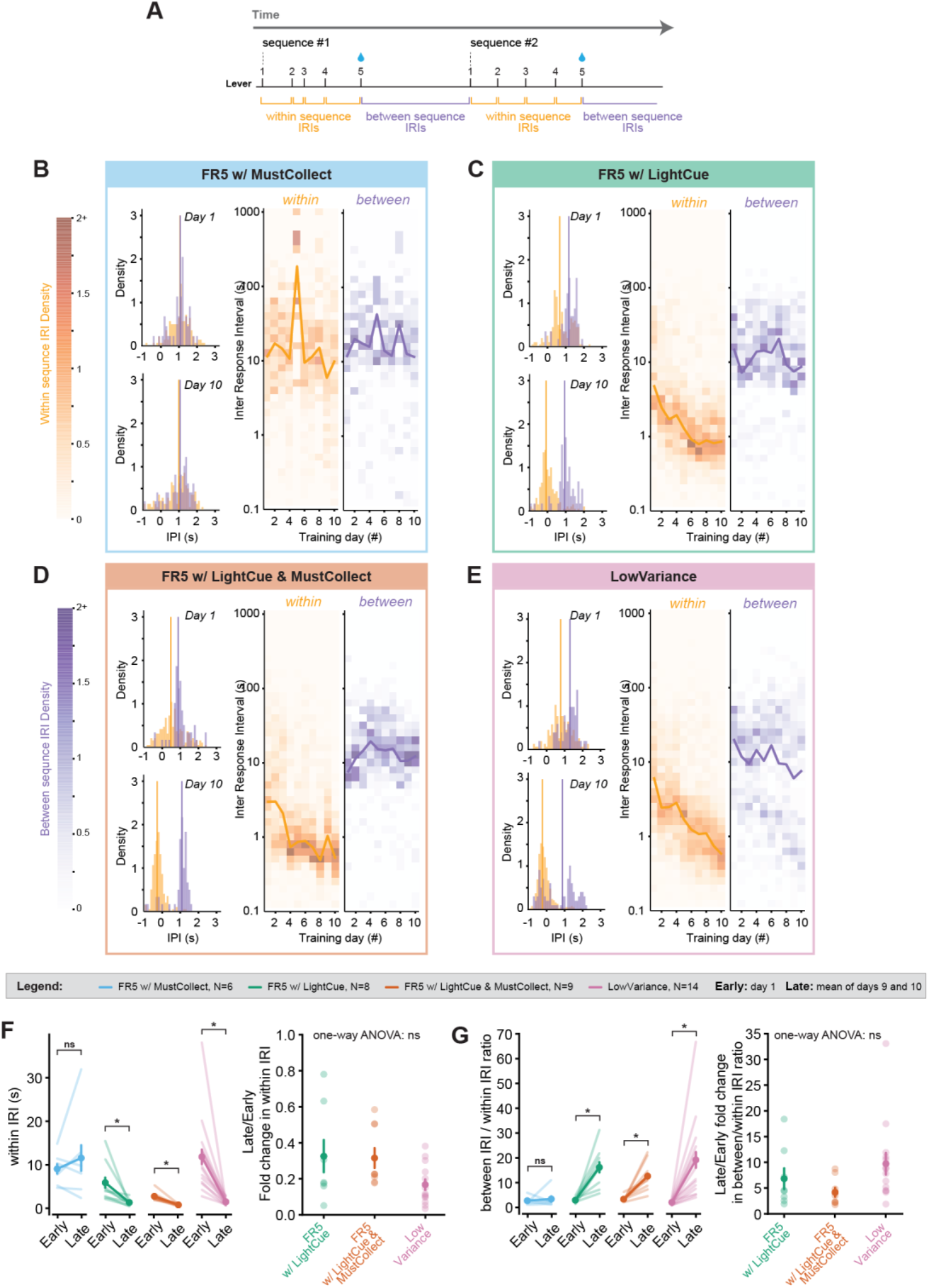
A light cue signaling sucrose delivery is necessary for clustering lever presses into bouts. **(A)** Diagram showing the distinction between inter-press intervals (IRIs) occurring within reinforced sequences and those occurring between sequences. **(B-E)** Plots showing the distribution of the two types of IRIs across training for example mice from each group. (left) Distribution of “within-sequence IRIs” and “between-sequence IRIs” on day 1 (top) and on day 10 (bottom). (center) Heatmap showing the distribution of “within-sequence IRIs” across days. Yellow line indicates the median “within-sequence IRI”. (right) Heatmap showing the distribution of “between-sequence IRIs” across days. Blue line indicates the median “between-sequence IRI”. **(F)** Comparison of “within-sequence IRIs” early (day 1) versus late (mean of days 9 and 10) for each group (left), and comparison of the late/early fold change for the groups with a significant difference. **(G)** Comparison of the ratio “between-sequence IRIs” / “within-sequence IRIs” early (day 1) versus late (mean of days 9 and 10) for each group (left), and comparison of the late/early fold change for the groups with a significant difference (right). Data presented as mean +/- S.E.M. * p < 0.05, ns, not significant.

### Direct reinforcement of low variance sequences does not produce more reproducible sequences than a traditional FR5

The distribution of “within-sequence IRIs” versus “between-sequence IRIs” provides a useful metric to quantify how well animals clustered their presses into bouts. However, it does not indicate whether sequences of presses become more reproducible across sessions. One goal of our experiment was to test the hypothesis that directly reinforcing low variance sequences is more effective than a traditional FR5 schedule in promoting reproducible behavior. To test this hypothesis, we used the variance of “within-sequence IRIs’’ as a quantitative metric to assess the reproducibility of the rhythm of presses within each sequence. While the variance of “within-sequence IRIs” is large because the rhythm is irregular early in training (**Figure 4A**), more stereotypical sequences later in training have lower “within-sequence IRI’’ variance (**Figure 4B**). To visualize how “within-sequence IRI” variance changed throughout training, we computed the IRI variance for each sequence and plotted heatmaps showing the distribution of these variances across sessions for example mice (**Figure 4C**). For each training strategy, we tested whether the “within-sequence IRI” variance changed from early (day 1) to late (mean of days 9 and 10) in training and found that the mean variance decreased only for mice trained on *FR5 w/ LightCue&MustCollect* and on *LowVariance* (**Figure 4D**, *FR5 w/ LightCue&MustCollect* - early: 11.9±1.4, late: 5.0±0.9, paired t-test p = 0.049; *FR5 w/ MustCollect* - early: 33.1±4.8, late: 48.7±10.6, paired t-test p = 0.18; *FR5 w/LightCue* - early: 18.5±3.8, late: 16.5±4.6, paired t-test p = 0.82; *LowVariance* - early: 26.3±5.4, late: 4.4±0.68, paired t-test p = 0.015). The late/early fold changes were not different between these two groups (**Figure 4E,** *FR5 w/LightCue&MustCollect*: 0.66±0.26; *LowVariance*: 0.35±0.086; paired t-test p = 0.23). These results show that the sequences of actions produced by mice trained on FR5 with a light cue that signals sucrose delivery and a reinforcer collection condition become more reproducible across training and that directly reinforcing reproducible sequences does not generate sequences that are less variable than those developed naturally.

**Figure 4:**
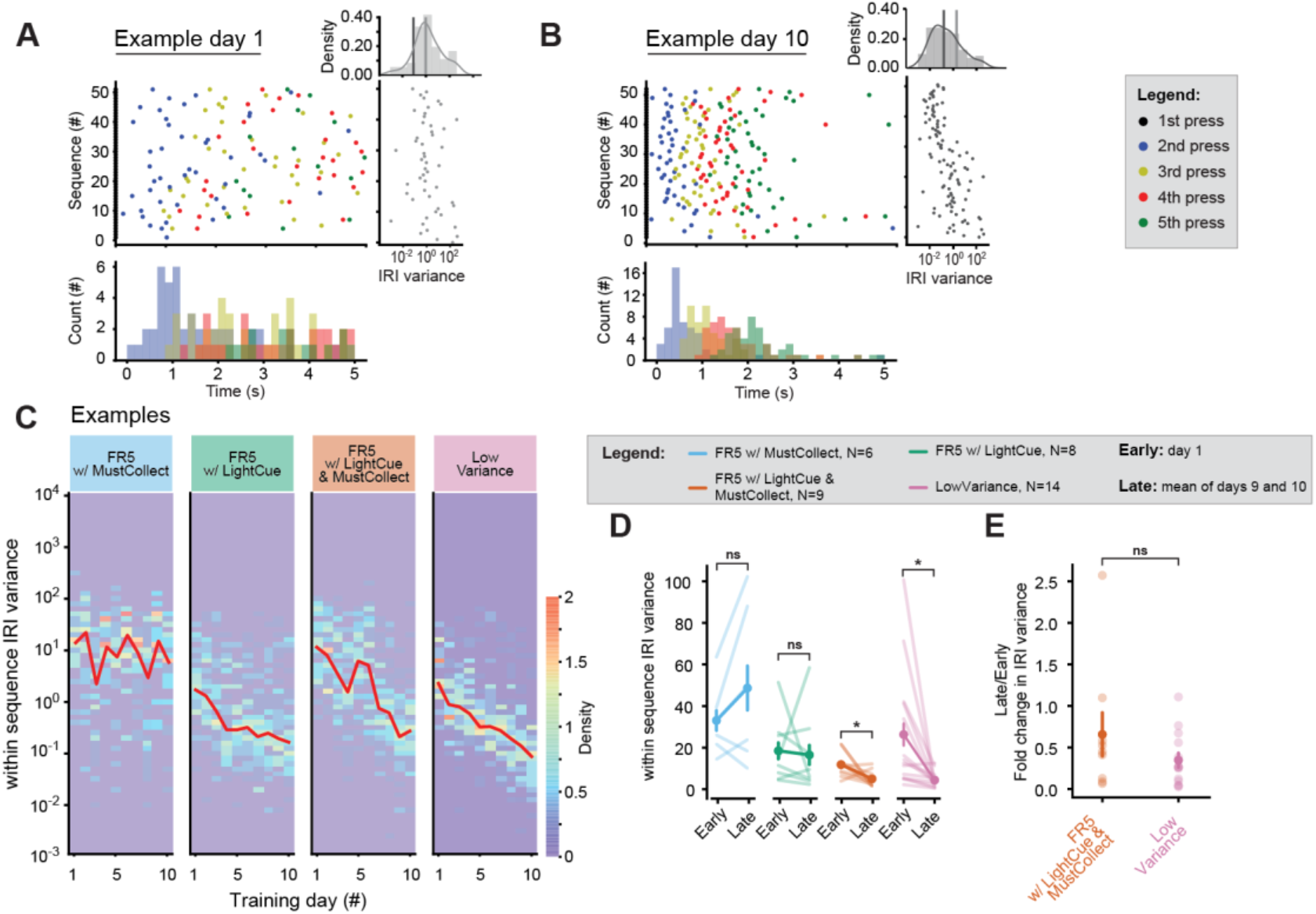
Direct reinforcement of low variance sequences does not produce more homogeneous sequences than an FR5 schedule. **(A-B)** Example raster plots (center) and post press time histograms (bottom) showing early **(A)** and late **(B)** sequences of presses aligned to the first press. The IRI variance for each sequence is plotted on the right panel and the distribution of these variances is shown in the top right plots. The light gray line shows the early variance median and the dark gray line shows the late variance median. **(C)** Heatmaps showing the distribution of “within-sequence IRI” variances for example mice from each group. **(D)** Comparison of the “within-sequence IRI” variance of reinforced sequences between early and late in training for each group. **(E)** Comparison of the late/early fold change for the groups with a significant difference. Data presented as mean +/- S.E.M. * p < 0.05, ns, not significant.

### Individual lever presses become less variable with training

The rhythm of presses is one way to quantify the reproducibility of sequential behavior. Another metric is the detailed kinematics of the movement executed by a mouse each time it presses the lever. Using magnetic sensors, we measured the displacement of the lever during behavioral sessions and tested the hypothesis that individual lever pressing movements become more reproducible as training progresses. Examples of presses in a sequence midway through training (**Figure 5A**, left, day 5) and late in training (**Figure 5A**, right, day 10) illustrate how individual presses indeed became more reproducible. The progression in press reproducibility was also apparent when we computed the pairwise Pearson correlation coefficient between all presses across training (**Figure 5B**). To quantify this progression and specifically test whether correlations improved with time and whether there was a difference between mice trained on *FR5 w/ LightCue&MustCollect* and mice trained on *LowVariance*, we compared the mean pairwise correlation coefficients across groups and between midway through training and late in training (**Figure 5C**). We found that indeed, the pairwise correlation coefficient between presses increased from midway to late in training and that mice trained on *FR5 w/ LightCue&MustCollect* had more reproducible presses than mice trained on the *LowVariance* paradigm (*LowVariance*-mid: 0.41±0.029, *LowVariance-late*: 0.46±0.037, *FR5 w/LightCue&MustCollect*-early: 0.55±0.065, *FR5 w/LightCue&MustCollect*-late: 0.61±0.041; mixed ANOVA: group (between subject) F = 6.2, p = 0.027; time (within subject) F = 8.5, p = 0.012; Post-hoc t-tests: *FR5 w/ LightCue&MustCollect_vs_LowVariance* p = 0.047, Mid_vs_Late p = 0.0096). These results indicate that mice produce presses that are less variable as they progress through training and confirm that the *LowVariance* training strategy did not produce more homogeneous actions.

**Figure 5:**
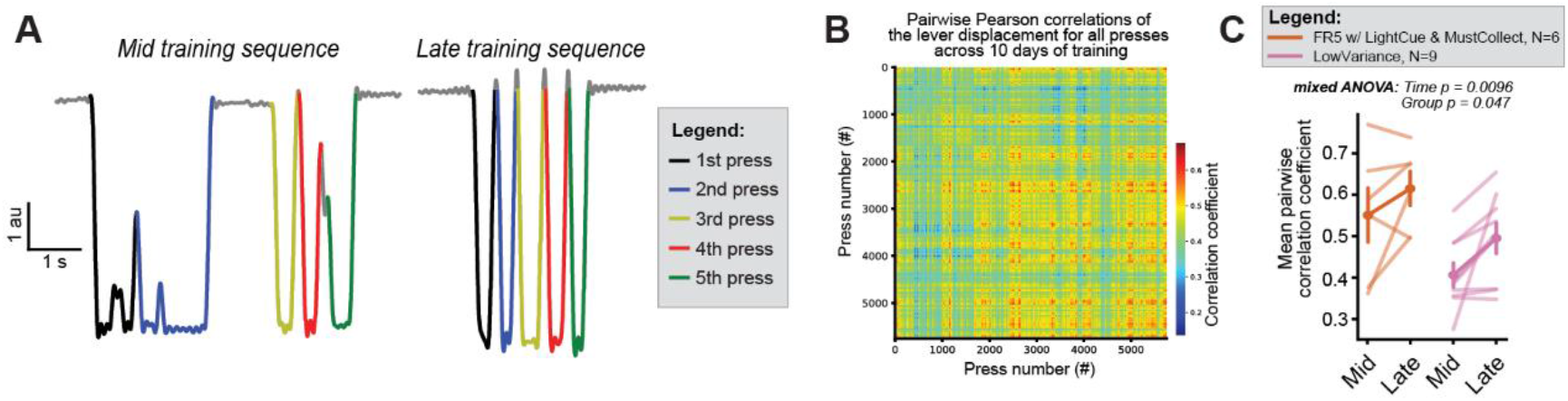
Individual presses become less variable with training. **(A)** Example sequences of five presses midway (day 5) through training (left) and late (day 10) in training (right) from one mouse. **(B)** Heatmap showing the pairwise correlation coefficients for the lever displacement of all presses of an example mouse trained on *LowVariance*. Correlations increase with time, showing presses become more similar to each other. **(C)** Comparison of the pairwise correlation coefficients across time and between *FR5 w/LightCue&MustCollect* and *LowVariance* groups. Data presented as mean +/- S.E.M.

### The *LowVariance* strategy induces a decrease in within-sequence inter response intervals but fails to produce sequences of five lever presses

Mice trained using the LowVariance strategy learned to press the lever (**Figure 1**), gradually increased their rate of pressing (**Figure 2**), clustered their presses (**Figure 3**) and reduced the variance of their within-sequence inter-response intervals (**Figure 4**). However, this paradigm was designed to only reinforce a sequence when the “within-sequence IRI” variance was below a threshold, which resulted in a subset of “failed sequences” - those for which the IRI variance did not dip below the threshold (**Figure 6A**). For those animals, the IRIs between sequences are therefore composed of two distinct types of IRIs; those that occur between the last press of a “failed” sequence and the first press of the next attempted sequence (**Figure 6A**, red “post failed sequence IRIs”) and those that occur between the last press of a reinforced sequence and the first press of the subsequent sequence (**Figure 6A**, blue “post reinforced sequence IRIs”). The bimodal distribution of “between-sequence IRIs” observed in the example mouse shown in Figure 3E suggests that these two types of IRIs change differentially with training. To test this possibility, we separated each type and compared IRIs early versus late in training. We found that, in addition to the “within attempted sequence IRIs”, the “post failed sequence IRIs” were dramatically shortened while the “post rewarded sequence IRIs” remained unchanged (**Figure 6B**, Within attempted sequence IRIs - early: 9.8±0.96 s, late: 1.5±0.16 s, paired t-test p = 3.7e-5; Post reinforced sequence IRIs - early: 18.3±1.7 s, late: 20.7±2.4 s, paired t-test p = 0.46; Post failed sequence IRIs - early: 16.9±2.2 s, late: 1.4±0.15 s, paired t-test p = 3.8e^-4^). Importantly, the late/early ratio was not different between “within attempted sequence IRIs” and “post failed sequence IRIs” (**Figure 6C**, Within attempted sequence IRIs: 0.17±0.029; Post failed sequence IRIs: 0.11±0.024; paired t-test p = 0.16), suggesting that while *LowVariance* mice successfully generated clustered lever presses, they failed to generate sequences of 5 presses.

**Figure 6:**
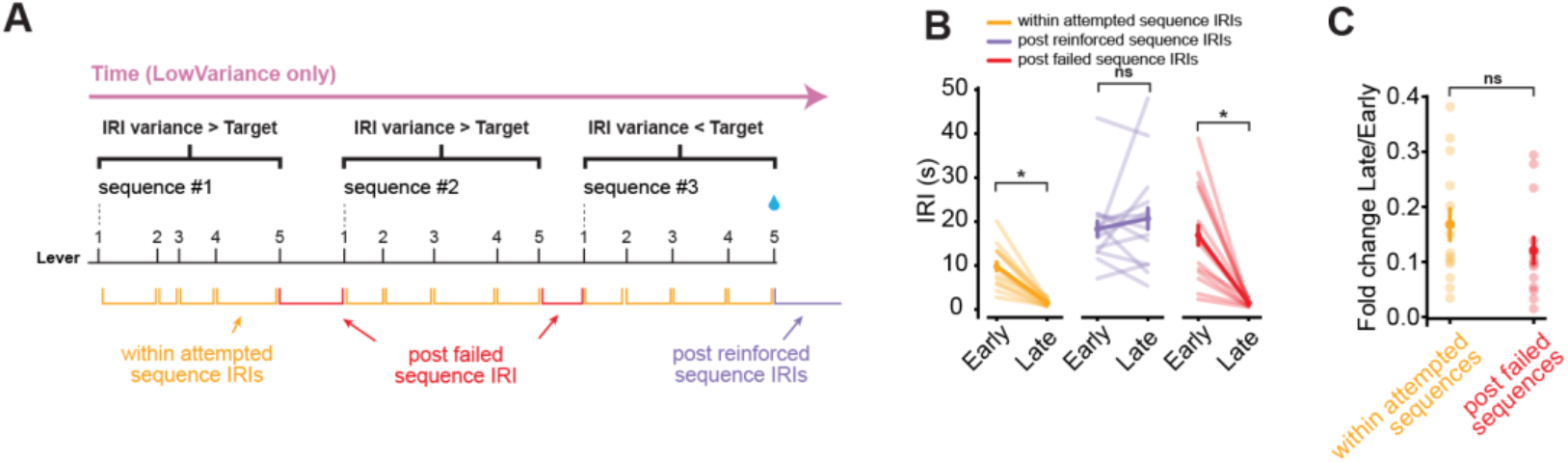
The LowVariance strategy induces a decrease of within-sequence interresponse intervals but fails to produce sequences of five lever presses. **(A)** Diagram showing the different types of IRIs for mice performing the *LowVariance* task. **(B)** Comparison of the early vs late IRIs for each type of IRI in *LowVariance* mice. **(C)** Comparison of the late/early fold change for each type of IRI that was significantly different. Data presented as mean +/- S.E.M. * p < 0.05, ns, not significant.

## Discussion

Automatic stereotypical behaviors are evolutionarily advantageous but can become maladaptive in disease states, such as OCD (Burguière et al., 2015) or SUD (Pierce and Vanderschuren, 2010). Studying these behaviors in controlled laboratory settings requires an understanding of the experimental conditions that promote the emergence of automaticity and reproducibility. Here, we sought to test whether specific schedules of reinforcement and contingencies within operant conditioning tasks were effective at promoting the generation of highly reproducible sequences of actions. Using several variations of an FR5 lever pressing task, we found that the presence of a light cue signaling sucrose delivery was a necessary component both for learning to press the lever (**Figure 1**) and for clustering presses into bouts (**Figures 2–4**), while a reinforcer collection condition did not dramatically improve sequencing behavior. In addition, we tested whether directly reinforcing low-variance sequences of lever presses (*LowVariance*) was more effective than a traditional FR5 strategy at generating reproducible actions and found that, while it effectively promoted an increase in response rate, the strategy we tested failed to generate predefined sequences of actions (ie. 5 lever presses) and to produce lever presses whose kinematics were less variable than those produced during FR5 training.

One striking result from our study is the learning deficits observed in the *FR5 w/ MustCollect* group, which was the only group without a cue that signaled sucrose delivery. Not only did these animals not learn to cluster their lever presses as well as the other groups (**Figures 2–4**), but they also already showed deficits during the acquisition of FR1 compared to the *FR5 w/LightCue&MustCollect* group (**Figure 1**). This finding is consistent with previous studies showing that “signaled” reinforcers reinforce behavior more effectively than “unsignaled” reinforcers (Branch, 1977; Doughty and Lattal, 2003; Lewis et al., 1974; Marcucella and Margolius, 1978; Sanderson et al., 2014; Schachtman and Reed, 1992) and that temporal proximity of reinforcement increases reinforcement learning efficiency (Arbel et al., 2017; Foerde and Shohamy, 2011; Peterburs et al., 2016; Weinberg et al., 2012; Yin et al., 2018). Indeed, as the reward cue provides an immediate proxy for sucrose availability, there is no need for the mouse to check whether a reinforcer was delivered and the duration for which action completion must be maintained in short term memory is reduced to virtually zero (Foerde and Shohamy, 2011).

While the light cue was effective at indicating sucrose delivery and contributed to reinforcing lever pressing, “post failed sequence IRIs” in the *LowVariance* group were indistinguishable from “within attempted sequence IRIs”, suggesting the animals failed to generate sequences of 5 presses (**Figure 6**). This pattern of IRIs reveals that under this particular contingency, mice exclusively depended on the light cue to stop their bout of presses. Therefore, while the light cue may not have promoted animals generating only 5 presses per bout, it may have allowed them to cluster their lever presses without developing an internal representation of sequential behavior per se – i.e. counting the number of responses. One interpretation is that the *LowVariance* group simply engaged a stereotypical rhythm of presses and waited for the light cue to signal they should stop their ongoing bout rather than executed a motor program consisting of a sequence of 5 presses (Wymbs et al., 2012). In one case, the sequential behavior is a predefined chunk of movements and time, similar to a habit, while in the other, it is a repeating action motif that is halted by external factors, perhaps similar to seeking behavior. Our study shows that one can easily masquerade as the other in the presence of a “signaled” reinforcer, which should be considered carefully when analyzing sequence behaviors.

Generally, more valuable reinforcers are more effective at changing behavior than less valuable reinforcers in positive reinforcement settings (Baron et al., 1992; Blakely and Schlinger, 1988; Schlinger et al., 1990). In the *LowVariance* group, the reward rate falls between 0 and 1 reinforcer per 5 presses depending on the mouse’s performance, while for a mouse performing a traditional FR5 the reward rate is 1 reinforcer to 5 presses. This difference likely explains why *FR5 w/ LightCue&MustCollect* mice pressed the lever at an overall faster rate than *LowVariance* mice (**Figure 2C**). Interestingly, it also suggests that *LowVariance* mice were able to achieve mostly comparable performance compared to *FR5 w/LightCue&MustCollect* when looking at measures of reproducibility (**Figures 2–5**) despite operating under a less efficient reinforcement paradigm. This effect is interesting in light of habitual behaviors. Indeed, a schedule of reinforcement in which the contingency between reward and action is uncertain, such as a random interval schedule, produces behaviors that are habitual (as defined by their insensitivity to reward devaluation, (Dezfouli and Balleine, 2012)). It remains to be investigated whether these habitual behaviors are also more stereotypical with regards to action kinematics and whether the homogeneity of actions comprising habits allows them to become reinforced at a lesser cost than more variable behaviors.

Together, we showed the complex relationship between operant contingencies and the induction of reproducible lever pressing patterns. We demonstrated the importance of a cue signaling reinforcer delivery on this behavior, showing that this cue was effective at increasing learning rates but resulted in mice pressing the lever in fast succession until the cue turned on, rather than pressing it a fixed number of times. Finally, we showed that a training strategy that directly reinforced sequences with low variance intervals was not more effective than a traditional fixed ratio schedule at promoting reproducible action execution. Together, our findings provide insights into the parameters of behavioral training that promote reproducible behavioral sequences.

## Acknowledgements

This work was supported by NIH grants DA042111 and DA048931 to E.S.C., 5T32MH065215-18 to M.C., as well as by funds from Brain and Behavior Research Foundation, the Whitehall Foundation, and the Edward Mallinckrodt, Jr. Foundation to E.S.C.

